# Loss of SDHB reprograms energy metabolisms and inhibits high fat diet induced metabolic syndromes

**DOI:** 10.1101/259226

**Authors:** Chenglong Mu, Biao Ma, Chuanmei Zhang, Guangfeng Geng, Xinling Zhang, Linbo Chen, Meng Wang, Jie Li, Tian Zhao, Hongcheng Cheng, Qianping Zhang, Kaili Ma, Qian Luo, Rui Chang, Qiangqiang Liu, Hao Wu, Lei Liu, Xiaohui Wang, Jun Wang, Yong Zhang, Yungang Zhao, Li Wen, Quan Chen, Yushan Zhu

**Affiliations:** State Key Laboratory of Medicinal Chemical Biology, Tianjin Key Laboratory of Protein Science, College of Life Sciences, Nankai University, Tianjin 300071, China.; State Key Laboratory of Membrane Biology, Institute of Zoology, Chinese Academy of Sciences, Beijing 100101, China.; Tianjin Key Laboratory of Exercise and Physiology and Sports Medicine, Tianjin University of Sport, Tianjin 300381, China.

**Keywords:** Complex II, Energy metabolism, Mitochondria, Obesity, SDHB

## Abstract

Mitochondrial respiratory complex II utilizes succinate, key substrate of the Krebs cycle, for oxidative phosphorylation, which is essential for glucose metabolism. Mutations of complex II cause cancers and mitochondrial diseases, raising a critical question of the (patho-)physiological functions. To address the fundamental role of complex II in systemic energy metabolism, we specifically knockout SDHB in mice liver, a key complex II subunit that tethers the catalytic SDHA subunit and transfers the electrons to ubiquinone, and found that SHDB deficiency abolishes the assembly of complex II without affecting other respiration complexes while largely retaining SDHA stability. SHDB ablation reprograms energy metabolism and hyperactivates the glycolysis, Krebs cycle and β-oxidation pathways, leading to catastrophic energy deficit and early death. Strikingly, sucrose supplementation or high fat diet resumes both glucose and lipid metabolism and prevent early death. Also, SDHB deficient mice are completely resistant to high fat diet induced obesity. Our findings reveal that the unanticipated role of complex II orchestrating both lipid and glucose metabolisms, and suggest that SDHB is an ideal therapeutic target for combating obesity.

## Introduction

Mitochondria are cellular centers for ATP production via the Krebs cycle and oxidative phosphorylation (OXPHOS). They also serve as a central regulator for cell physiology, from the synthesis of intermediary metabolites and ROS production to modulation of cell death pathways. OXPHOS comprises four respiratory complexes, which can form a supercomplex depending on the energy status within a cell. Complex II, also named succinate dehydrogenase (SDH), is a key enzymatic complex that catalyzes the oxidation of succinate to fumarate in Krebs cycle. Complex II is a highly conserved heterotetrameric protein complex consisting of four nuclear-encoded subunits SDHA, SDHB, SDHC, and SDHD [1, 2]. SDHC and SDHD are embedded in the inner mitochondrial membrane, and the SDHC/SDHD dimer binds the peripheral membrane protein SDHB, which tethers the catalytic SDHA subunit so that it faces the mitochondrial matrix [3]. SDHA harbors a covalently bound FAD cofactor that is required for the oxidation of succinate [4]. The two electrons that result from succinate oxidation are channeled through the three iron-sulfur clusters in SDHB to ubiquinone [5]. Complex II is thus unique and functions at the crossroads of the Krebs cycle and OXPHOS pathways [6]. The genes encoding complex II subunits were found to be mutated in various types of tumor, such as paraganglioma (PGL) [7], pheochromocytoma (PHEO) [8], gastrointestinal stromal tumor (GIST) [9], renal cell carcinoma and Leigh syndrome [10]. Complex II is also reported to be a tumor suppressor [11–13]. Complex II is thus considered as a therapeutic target for highly debilitating pathologies including cancer, Parkinson’s disease [14] and inflammatory diseases, with potential clinical relevance. Ischemic accumulation of succinate via reversal of its SDH activity drives the mitochondrial ROS production upon reperfusion that underlies IR injury in a range of tissues[15]. Such accumulation of succinate also plays a role in inflammatory and hypoxic signaling [16].

Mitochondria are the principal sites for β-oxidation, which breaks down fatty acids for energy production. The final products of fatty acid β-oxidation are acetyl-CoA, NADH, and FADH2, which enter into Krebs cycle and OXPHOS for ATP production. The metabolism of fatty acid and glucose are tightly linked through gluconeogenesis and *de novo* lipid synthesis pathways in liver and other metabolic organs, depending on the feeding status. Under nutrient-rich conditions, glycolytic products are used to synthesize fatty acids in liver through *de novo* lipogenesis, and the lipids are stored in lipid droplets in fat tissue, or secreted into the circulation as VLDL particles. In the fasted state, the liver produces glucose through breakdown of glycogen and gluconeogenesis. Prolonged fasting promotes lipolysis in adipose tissue; the released fatty acids are converted into ketone bodies in the liver using acetyl-CoA produced through mitochondrial β oxidation [17]. The liver thus lies at the center of energy metabolism within our bodies and orchestrates the systemic metabolic balance of glucose, lipids, and amino acid. Aberrant energy metabolism in the liver causes metabolic syndromes including insulin resistance, diabetes, and nonalcoholic fatty liver diseases [18]. Mitochondria are highly enriched in liver tissue and play essential roles in tightly regulating these metabolic processes for homeostasis[19]. There is growing evidence that mitochondrial complex II is tightly associated with metabolic regulation at the molecular level, but less is known about its physiological significance. Here we show that complex II is an essential regulator of energy metabolism and plays important roles in orchestrating both glucose and fatty acid metabolism. To our surprise, liver-knockout SDHB mice are completely resistant to high fat diet induced obesity and showed complex II may be a candidate target for fighting obesity.

## Results

### Liver-specific knockout of SDHB shortens the lifespan in mice

To understand the critical role of complex II in mitochondrial respiration and regulating systemic energy metabolism, we created mice with knockout of SDHB, which is one subunit of complex II. The whole-body knockout of SDHB is embryonic lethal (Fig EV1A), which indicates that complex II is essential for development. Given the central role of liver in systemic energy metabolism, we generated a liver-specific knockout of SDHB in mice (Fig 1A-1C and Fig EV1B) (hereafter *Sdhb^-/-^*). These mice were viable at birth but died at about two months of age (Fig 1D). The 5-week-old *Sdhb^-/-^* mice were completely normal in body weight and daily activities (Fig EV2A and EV2B). However, the 8-week-old *Sdhb^-/-^* mice showed severe growth retardation, which started soon after weaning (Fig 1E and 1F). Liver tissue analysis showed that ablation of SDHB resulted in the loss of complex II activity as assayed by complex II staining (Fig 1I) and DCPIP test (Fig 1J), even though the stability of the catalytic subunit, SDHA, was largely maintained (Fig 1H). The assembly and stability of components of other OXPHOS complexes were unaffected (Fig 1G and 1H). Furthermore, oxygen consumption ratio (OCR) analysis revealed that mitochondria from *Sdhb^-/-^* liver tissues did not respond to succinate (Fig 1K), while the responses to malate and glutamate were unaffected (Fig 1L). These results indicate that the activity of complex II was specifically ablated in the liver of *Sdhb^-/-^* mice, while the activity of other complexes was maintained.

**Figure 1.**
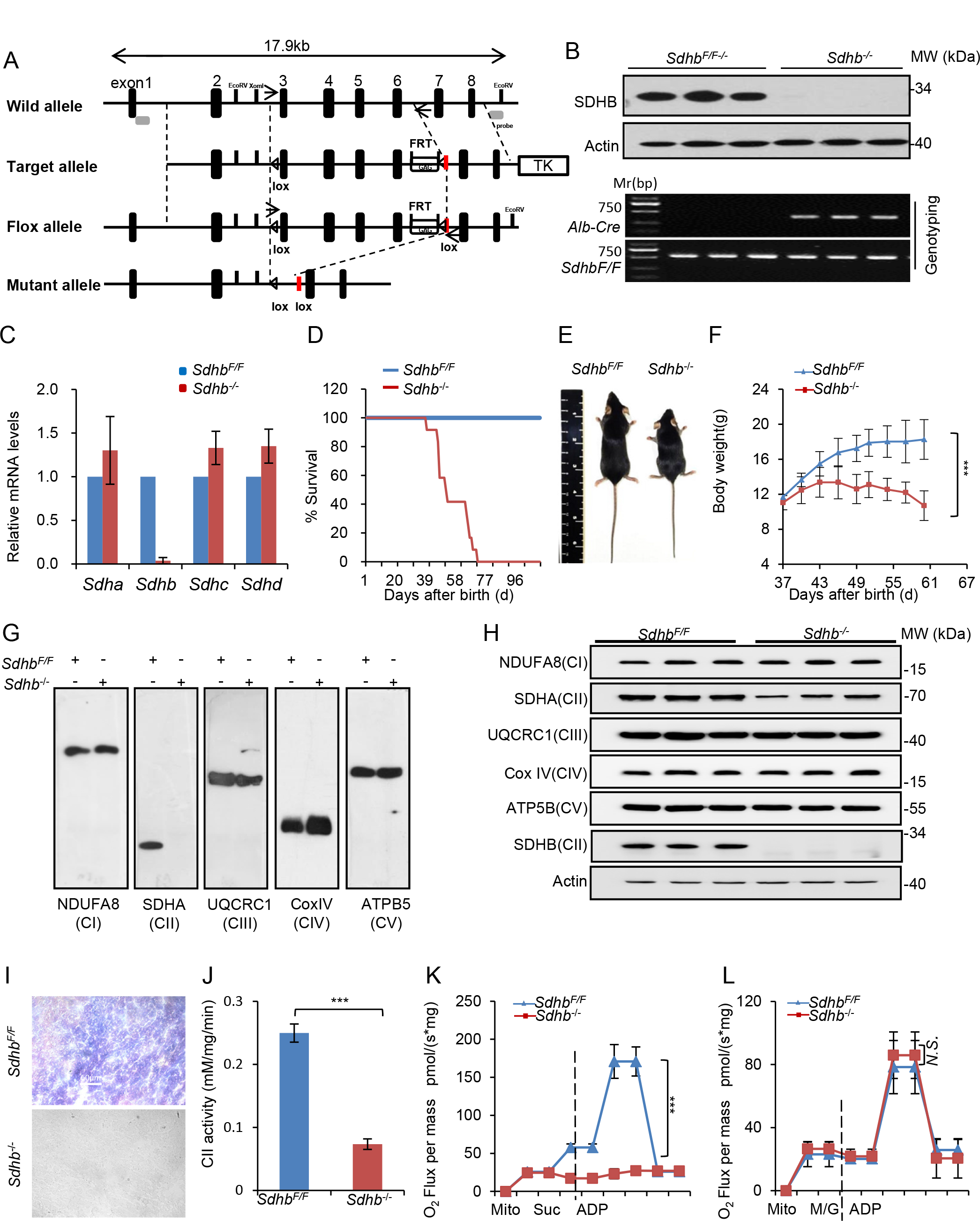
Construction and phenotypic analysis of SDHB liver knockout (*Sdhb^-/-^*) mice. A. Gene targeting strategy for *Sdhb^F/F^* mice. Lox sites were inserted before Exon 3 and after Exon 6 of the *Sdhb* gene. Exons 3, 4, 5 and 6 are deleted in the presence of Cre recombinase. B. Genotyping (lower panel) and SDHB protein levels in liver tissues (upper panel, ACTIN was used as the loading control.) of *Sdhb^F/F^* and *Sdhb*^-/-^ mice (n=3). C. Relative mRNA levels of the four subunits of complex II in *Sdhb^F/F^* and *Sdhb*^-/-^ mice (n=6). D. Survival curves of *Sdhb^F/F^* and *Sdhb*^-/-^ mice fed with a Chow diet (CD) (n=5-7), E. Body sizes of *Sdhb^F/F^* and *Sdhb*^-/-^ mice at 8 weeks of age. F. Body weight curve of *Sdhb*^-/-^ and *Sdhb^F/F^* mice (n=6). G. Analysis of the assembly of OXPHOS complexes I-V in the livers of *Sdhb^F/F^* and *Sdhb*^-/-^ mice by blue native PAGE and western blotting. The subunits analyzed are NDUFA8 for complex I (CI), SDHA for complex II (CII), UQCRC1 for complex III (CIII), CoxIV for complex IV (CIV), and ATP5B for complex V (CV), ACTIN was used as the loading control. H. Levels of representative subunits of OXPHOS (oxidative phosphorylation) complexes in the livers of *Sdhb^F/F^* and *Sdhb*^-/-^ mice (n=3). I. Representative images of SDH (succinate dehydrogenase) staining for complex II activity in the livers of *Sdhb^F//F^* and *Sdhb*^-/-^ mice (n=4). Scale bar: 50 μm. J. Mitochondrial complex II activity assay in liver from *Sdhb^F/F^* and *Sdhb*^-/-^ mice by the DCPIP method (n=3). K. Measurement of mitochondrial OCR (oxygen consumption ratio) in *Sdhb^F/F^* and *Sdhb*^-/-^ mice in response to the complex II substrate succinate (n=6). L. Measurement of mitochondrial OCR in *Sdhb^F/F^* and *Sdhb*^-/-^ mice in response to the complex I pathway substrates malate and glutamate (n=6). Bar graphs represent mean ± SEM, \**p*<0.05, ** *p*<0.01, *** *p*<0.001, *N.S*., no significance.

### Deficiency of complex II in liver accelerates glucose and fat consumption

Considering that complex II is an important component of Krebs cycle and OXPHOS for glucose metabolism, we first checked the serum glucose levels in 8-week-old mice. Intriguingly, the serum glucose level in *Sdhb^-/-^* mice was significantly lower than in *Sdhb*^F/F^ mice (Fig 2A). PAS (Periodic Acid-Schiff) staining also showed that *Sdhb^-/-^* mice had a lower level of glycogen in liver (Fig 2B and 2C), indicating that glucose consumption may be increased in *Sdhb^-/-^* mice. To further validate this, mice were injected with glucose at the dosage of 2 g/kg body weight, then dynamic serum glucose concentrations were monitored and liver tissues were analyzed by PAS staining. The rate of glucose consumption was indeed higher in *Sdhb^-/-^* mice by glucose tolerance test (GTT) (Fig 2D), while glycogen accumulation in *Sdhb^-/-^* mice was significantly inhibited (Fig 2E and 2F). It was noticed that the levels of serum glucose and liver glycogen of 5-week-old *Sdhb^-/-^* mice were identical with those of *Sdhb*^F/F^ controls (Fig EV2E and EV2F). Taking these data together, we conclude that glucose catabolism is accelerated by the loss of complex II activities in 8-week-old mouse.

**Figure 2.**
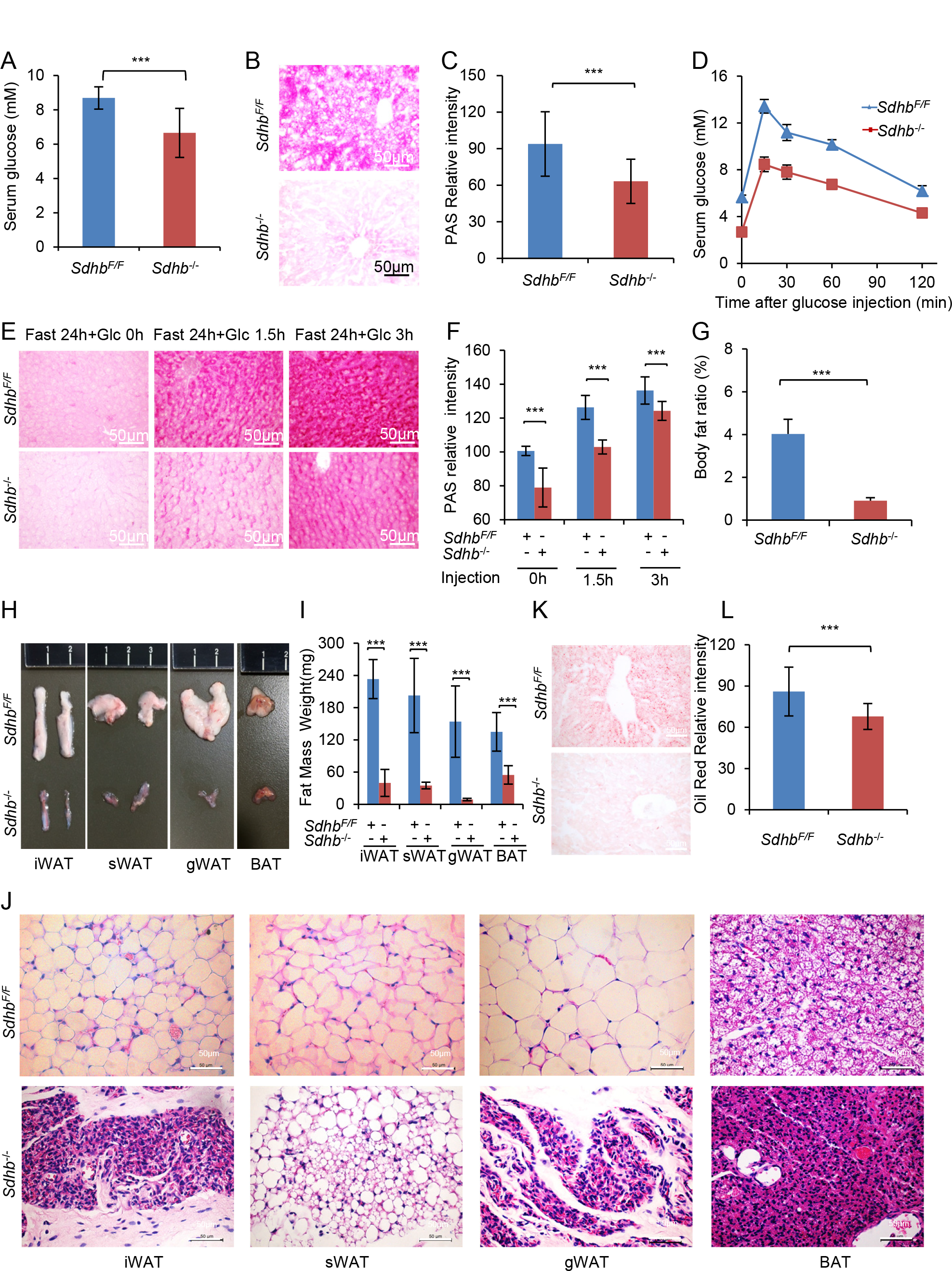
Loss of SDHB in liver increases the catabolism of glucose and fatty acids. A. Serum glucose levels in *Sdhb*^-/-^ and *Sdhb^F/F^* mice (n=5). B. Representative images of PAS (Periodic Acid-Schiff) staining for glycogen in the liver tissue of *Sdhb*^-/-^ and *Sdhb^F/F^* mice. C. Quantification of the intensity of PAS staining in B (n=4; at least ten microscope fields were examined for each liver section). D. Glucose tolerance test (GTT) in *Sdhb*^-/-^ and *Sdhb^F/F^* mice at 8 weeks of age (n=5). Bar graphs represent mean ± SEM. E. Representative images of PAS staining for glycogen accumulation in liver tissue of *Sdhb^F/F^* and *Sdhb*^-/-^ mice at 1.5 hours or 3 hours after 20% glucose intraperitoneal injection. Scale bar: 50 μm. F. Quantification of the intensity of PAS staining in E (n=3; 10 microscope fields per liver section). G. Body fat ratio of *Sdhb*^-/-^ and *Sdhb^F/F^* mice at 8 weeks of age (n=5). H. Images of the main white adipose tissues (WAT), including inguinal WAT (iWAT), scapular WAT (sWAT) and gonadal WAT (gWAT), and brown adipose tissue (BAT) between the scapulae of *Sdhb*^-/-^ and *Sdhb^F/F^* mice. I. Quantification of the adipose tissue mass in H (n=5). J. Representative sizes of adipocytes in iWAT, sWAT, gWAT and BAT of *Sdhb^F/F^* and *Sdhb*^-/-^ mice respectively, H&E staining of paraffin sections (n=4). Scale bar: 50 μm. K. Representative images of Oil Red O-stained lipid droplets in the liver of *Sdhb^F/F^* and *Sdhb*^-/-^ mice. Scale bar: 50μm. L. Quantification of the intensity of Oil Red O staining in H (n=4; 10 microscope fields per liver section). Bar graphs represent mean ± SEM, \**p*<0.05, ** *p*<0.01, *** *p*<0.001, *N.S*., no significance.

In addition to the drastic changes in glucose consumption in 8-week-old *Sdhb^-/-^* mice, we also observed that there was a significant decrease in the body fat ratio (Fig 2G). Anatomic analysis showed that the masses of all white fat tissues (WAT), such as inguinal WAT (iWAT), scapular WAT (sWAT) and gonadal WAT (gWAT), and of brown adipose tissue (BAT), were dramatically decreased in *Sdhb^-/-^* mice (Fig 2H and 2I). Furthermore, the adipocytes in all WAT and BAT tissues were reduced in size (Fig 2J). Considering that the body fat ratio, adipose tissue mass, adipocyte size, and the staining of liver lipid droplets are identical in *Sdhb^-/-^* and control mice at 5 weeks of age (Fig EV2H, EV2I and EV2J), the decrease in fat tissue and adipocytes at 8 weeks of age may be due to increased lipid mobilization. Thus we checked the liver lipid levels in 8-week-old *Sdhb^-/-^* mice. Surprisingly, the levels of lipids were significantly reduced in the *Sdhb^-/-^* liver (Fig 2K and 2L), suggesting that fatty acid catabolism may be also accelerated in liver.

### Deficiency of SDHB in liver upregulates pathways involved in glucose and fat catabolism

To understand the molecular basis of these metabolic alterations in *Sdhb^-/-^* liver, we performed RNAseq analysis using livers from 8-week-old mice. The results showed that multiple genes involved in glycolysis, Krebs cycle and β-oxidation were upregulated in SDHB-deficient liver tissues (Fig 3A). Western blotting analysis of liver samples from 8-week-old mice further confirmed that enzymes involved in glucose transportation, Krebs cycle, fatty acid β-oxidation and the gluconeogenesis pathway (Fig 3B, 3C and 3D) were simultaneously upregulated. It is apparent that SDHB deficiency induces metabolic reprogramming which results in the enhanced catabolism of glucose and fats in *Sdhb^-/-^* tissues (Fig 3E). It is surprising that one key glycolytic enzyme, GCK, is downregulated at 8 weeks of age, as this is not in line with other results indicating accelerated glucose catabolism, such as lower serum glucose level (Fig 2A and 2D) and liver glycogen accumulation (Fig 2B, 2C, 2E and 2F). To account for this discrepancy, we reasoned that the feeding status of the mice may affect the glycolytic pathway. Thus we examined the protein levels of the main glycolytic enzymes in the liver tissue of 5-week-old mice, at which time the serum glucose levels were normal in *Sdhb^-/-^* mice (Fig EV2C). Intriguingly, the levels of glycolytic enzymes were increased in *Sdhb^-/-^* mice at 5 weeks of age, and proteins involved in glucose transportation, Krebs cycle and fatty acid β-oxidation were also upregulated (Fig EV2K). It seems that the decrease in serum glucose levels affects the liver glycolysis levels in *Sdhb*^-/-^ mice at 8 weeks of age. HIF1A is reported as a regulator of glucose metabolism in SDHB-deficient cells [20, 21]. Here we also found that HIF1A levels were high in *Sdhb*^-/-^ mice at both 5 and 8 weeks of age (Fig 2A and Fig EV2K). These data indicate that HIF1A may play some role in the metabolic reprogramming in *Sdhb*^-/-^ mice. Taking the above data together, we conclude that loss of complex II can upregulate glucose and fatty acid catabolic pathways.

**Fig 3.**
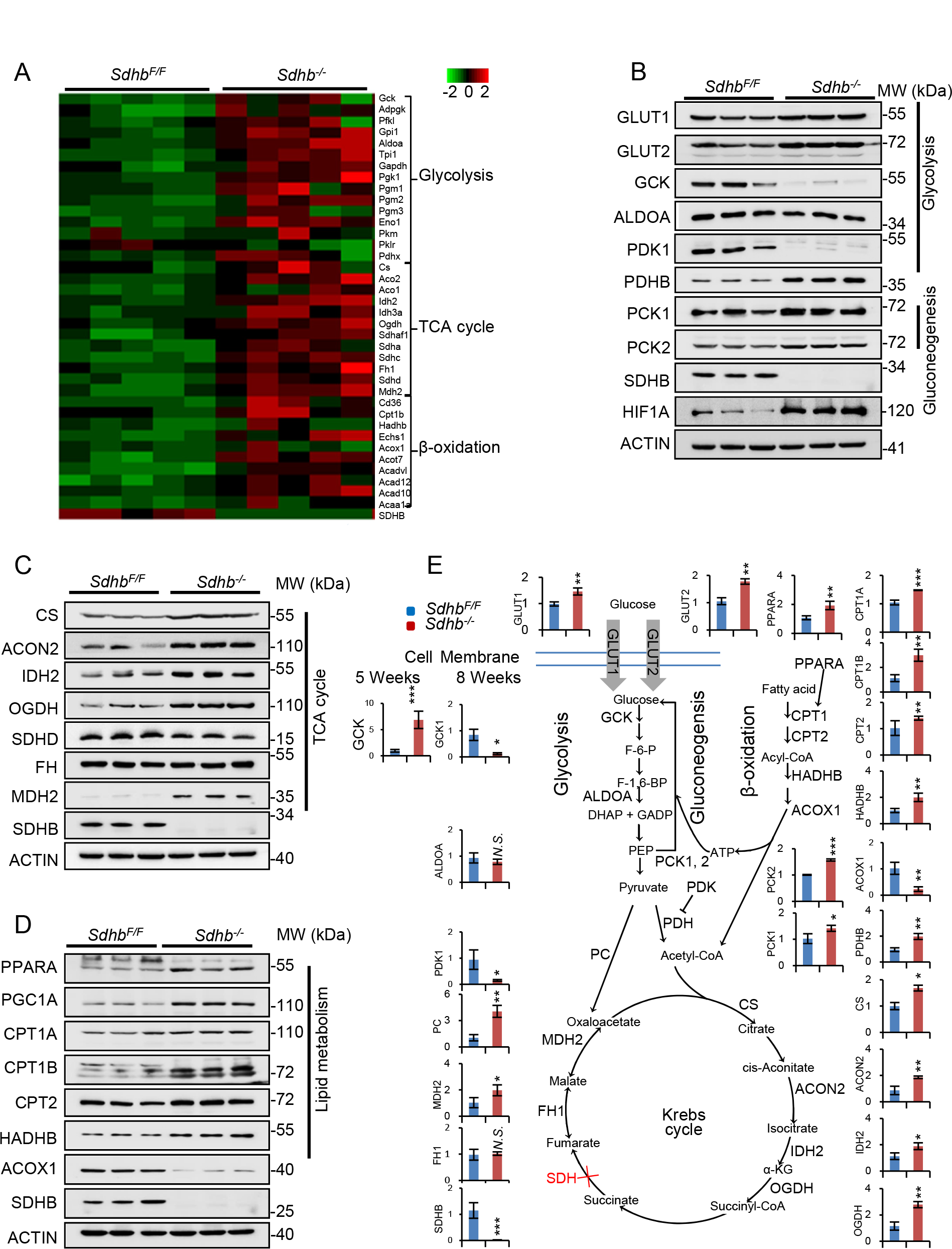
Loss of SDHB in liver upregulates glucose and fatty acid catabolism pathways. A. Heat map for RNA sequencing results of mRNAs involved in glycolysis, Krebs cycle and β-oxidation pathways in *Sdhb^F/F^* and *Sdhb*^-/-^ mice at 8 weeks of age (n=5). B. Levels of proteins involved in glucose transportation (GLUT1, GLUT2), glycolysis (GCK, PFKL, ALDOA, PDK1, PDHB) and gluconeogenesis (PCK1, PCK2) in liver tissue of *Sdhb^F/F^* and *Sdhb*^-/-^ mice at 8 weeks of age (n=3). ACTIN was used as the loading control. C. Protein levels of key Krebs cycle enzymes (CS, ACON2, IDH2, OGDH, SDHD, FH, MDH2 and SDHB) in the livers of *Sdhb^F/F^* and *Sdhb*^-/-^ mice at 8 weeks of age (n=3). ACTIN was used as the loading control. D. Protein levels of key enzymes in lipid metabolism (PPARA, PGC1A, CPT1A, CPT1B, CPT2, HADHB and ACOX1) in the livers of *Sdhb^F/F^* and *Sdhb*^-/-^ mice at 8 weeks of age (n=3). ACTIN was the loading control. E. Metabolic reprogramming profiles and quantification of the western blot results from B, C and D (n=3). Bar graphs represent mean ± SEM, \**p*<0.05, ** *p*<0.01, *** *p*<0.001, *N.S*., no significance.

### High levels of dietary carbohydrate can fully restore glucose and fatty acid metabolism in *Sdhb^-/-^* mice

We further speculated that energy exhaustion due to enhanced catabolism may cause the death of *Sdhb*^-/-^ mice. If so, supplementing the diet with energy sources may rescue these phenotypes. Carbohydrates and fats are the two most important energy substrates. Hence, we first supplemented the drinking water with 20% sucrose. As expected, addition of sucrose completely rescued the lifespan (Fig 4A) and growth retardation (Fig 4B and 4C) of *Sdhb*^-/-^ mice, and levels of blood glucose were restored to the normal levels (Fig 4D). Anatomic analysis further showed that the body fat ratio (Fig 4E), fat mass, and adipocyte size of WAT and BAT recovered to normal levels (Fig 4F, 4G and 4H). Liver fat accumulation was also increased in *Sdhb*^-/-^ tissues (Fig 4I and 4J). We further analyzed the protein levels of enzymes involved glucose transportation, glycolysis, Krebs cycle, β-oxidation, gluconeogenesis and HIF1A. Under sucrose treatment, key enzymes in glycolysis and some enzymes in Krebs cycle downstream of complex II were upregulated as well as HIF1A, while enzymes involved in β-oxidation, gluconeogenesis and Krebs cycle upstream of complex II were decreased (**Fig 4K**). These results indicated that when the body has a sufficient supply of carbohydrates, metabolism switches to a glycolysis-dependent pathway. Taking these data together, we conclude that supplementation with sucrose can meet the energy expenditure in *Sdhb*^-/-^ mice for survival, and carbohydrate metabolism can compensate for the greatly increased fatty acid catabolism in these animals.

**Figure 4.**
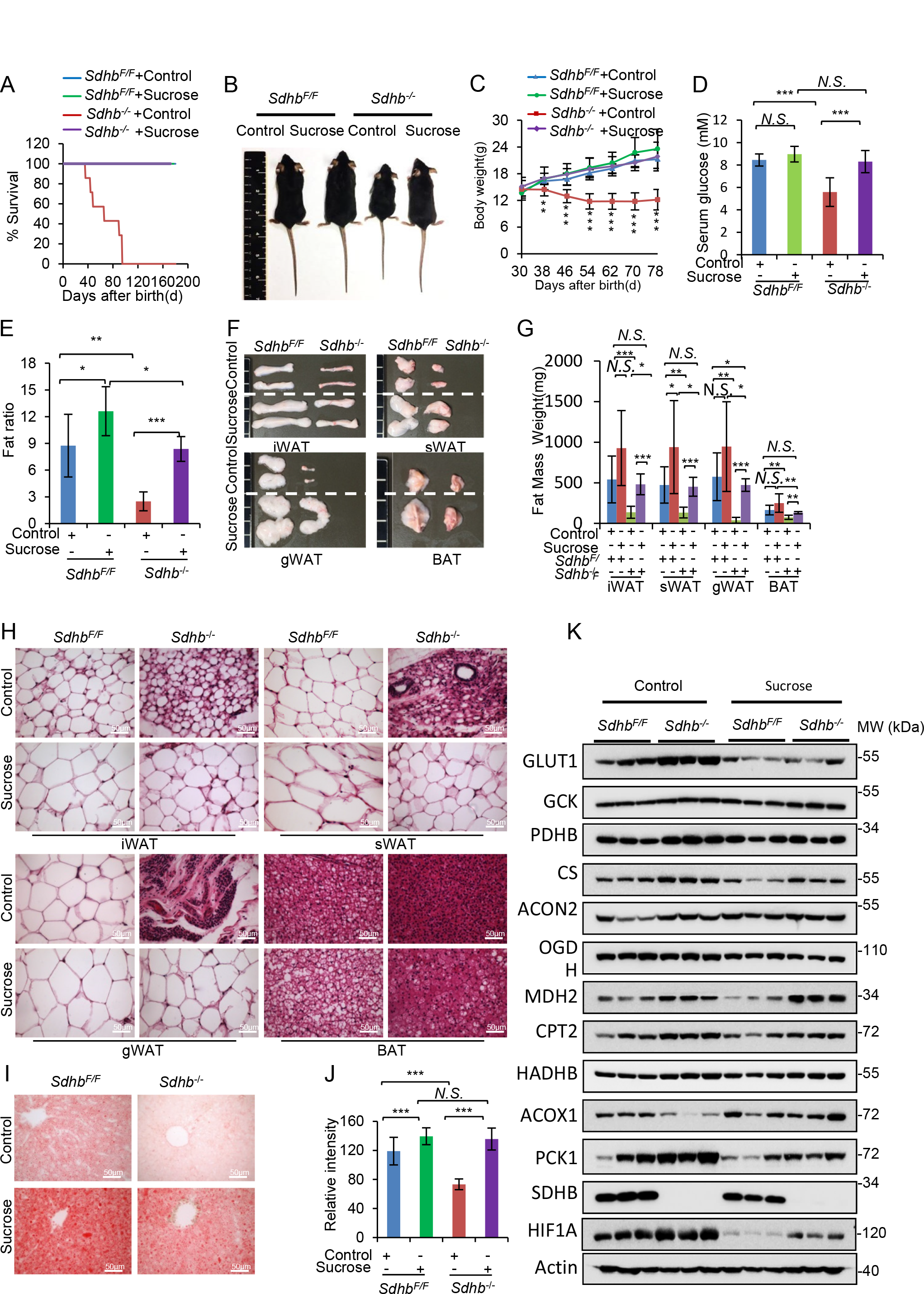
High sucrose treatment rescues the phenotypes of *Sdhb^-/-^* mice. A Survival curves of *Sdhb^F/F^* and *Sdhb*^-/-^ mice at 4 weeks of age treated with sucrose solution or water (control) for four weeks (n=6). B-E. Body sizes (B), body weight curves (C), serum glucose levels (D), and body fat ratios (E) in mice as described in A. F. Images of the main adipose tissues (iWAT, sWAT, gWAT, and BAT) from mice as described in A. G. Quantification of the masses of the adipose tissues shown in F (n=6; 6 microscope field for per adipose sections). H. Representative images of adipocytes in iWAT, sWAT, gWAT and BAT. Scale bar: 50 μm. I. Representative images showing Oil Red O staining of liver lipid droplets in mice as described in A. Scale bar: 50 μm. J. Quantification of the intensity of Oil Red O staining in I (n=5; 4 microscope fields were examined per liver section). K. Protein levels of enzymes involved in glycolysis (GLUT1, GCK, PKLR, PDHB), Krebs cycle (CS, ACON2, OGDH and MDH2), lipid metabolism (CPT2, HADHB and ACOX1), gluconeogenesis (PCK1) and HIF1A in the liver tissue of mice as described in A (n=3). Bar graphs represent mean ± SEM, \**p*<0.05, ** *p*<0.01, *** *p*<0.001, *N.S*., no significance.

### HFD restores glucose metabolism and lifespan in *Sdhb^-/-^* mice

Fatty acids are the most energy-abundant material within the body, so we speculated that high-fat diet (HFD) would compensate for the accelerated glucose metabolism in *Sdhb*^-/-^ mice. When *Sdhb*^-/-^ mice were given one month of HFD treatment beginning at 4 weeks of age, their body weights (Fig 5A and 5B) were fully restored. The HFD also led to recovery of the levels of blood glucose (Fig 5C) and liver glycogen (Fig 5D and 5E), and the body fat ratio (Fig 5G). Along with improved serum glucose levels, the glucose intolerance of *Sdhb*^-/-^ mice was decreased, even though glucose consumption was still higher than in HFD-treated *Sdhb*^F/F^ mice (Fig 5F). Moreover, HFD-treated *Sdhb*^-/-^ mice had identical adipose tissue mass (Fig 5H and 5I), adipocyte size (Fig 5J) and levels of liver fatty acid droplets (Fig 5K and 5L) as normal chow diet (CD)-fed mice. Lipid droplet levels in HFD-treated *Sdhb*^-/-^ mice were also lower than HFD-treated *Sdhb*^F/F^ mice, verifying the increased fatty acid catabolism in *Sdhb*^-/-^ mice. Furthermore, western blotting analysis showed that glycolysis enzymes were all decreased, while enzymes in Krebs cycle, β-oxidation and gluconeogenesis were still increased, in *Sdhb*^-/-^ mice when fed a HFD (Fig 5M). These results indicated that HFD-treated mice switched their metabolic pattern to the β-oxidation pathway, and serum glucose was maintained by gluconeogenesis. Taking these results together, we can conclude that complex II deficiency increases catabolism of energy substrates, and energy status can reprogram the metabolic pathways.

**Figure 5.**
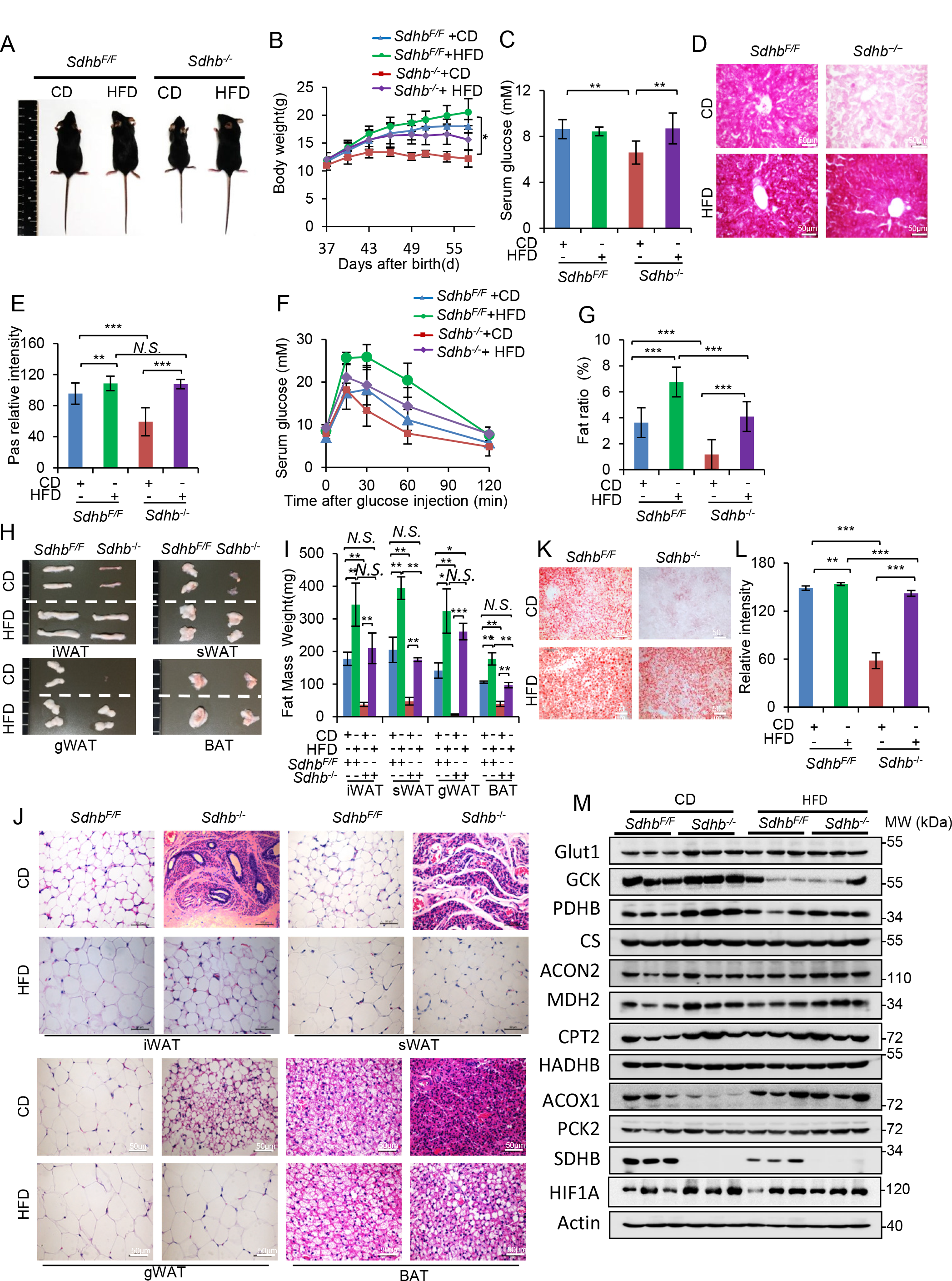
The phenotypes of *Sdhb^-/-^* mice are also rescued by HFD (high-fat diet). A-D Body sizes (A), body weight curves (B), serum glucose levels (C) and PAS staining for glycogen in liver tissues (D) from 4-week-old mice treated with HFD or CD for four weeks (n=6). E. Quantification of the intensity of PAS staining in D (n=4; 10 microscope fields per liver section). F. GTT assay in *Sdhb^F/F^* and *Sdhb*^-/-^ mice as described in A (n=6). G. Body fat ratio of *Sdhb^F/F^* and *Sdhb*^-/-^ mice as described in A (n=6). H. Images of the main adipose tissues (iWAT, sWAT, gWAT and BAT) from *Sdhb^F/F^* and *Sdhb*^-/-^ mice as described in A. I. Quantification of the masses of the adipose tissues shown in H (n=6). J. H&E staining of paraffin sections of iWAT, sWAT, gWAT and BAT tissue from H. Scale bar: 50 μm. K. Oil Red O staining for lipid droplet accumulation in the livers of mice as described in A. Scale bar: 50 μm. L. Quantification of the relative intensity of Oil Red O staining in K (n=4; 5 microscope fields per liver section). M. Protein levels of glucose transporter GLUT1, and enzymes involved in glycolysis (GCK, and PDHB), Krebs cycle (CS, ACON2, MDH2), lipid metabolism (CPT2, HADHB, ACOX1), gluconeogenesis (PCK1) and HIF1A in *Sdhb^F/F^* and *Sdhb*^-/-^ mice treated with HFD or CD (n=3). Bar graphs represent mean ± SEM, \**p*<0.05, ** *p*<0.01, *** *p*<0.001, *N.S*., no significance.

### The restoration of *Sdhb^-/-^* phenotypes by HFD can be inhibited by Etomoxir

Etomoxir (ETO) is an inhibitor of β-oxidation which acts by inhibiting carnitine palmitoyltransferase I (CPT1) [22]. To examine whether fatty acid β-oxidation within mitochondria provides the vital energy sources for rescuing the phenotype of *Sdhb*^-/-^ mice, we treated these mice with ETO under HFD conditions. The results showed that two-week ETO treatment significantly reduced the body weight (Fig 6A and 6B) and serum glucose levels of HFD-treated *Sdhb*^-/-^ mice (Fig 6C). Moreover, ETO treatment also dramatically inhibited fat consumption in *Sdhb*^-/-^ liver tissues, so that the level of liver fat droplets was identical level to that in *Sdhb*^F/F^ mice (Fig 6D and 6E). This indicates that the process of HFD-induced energy compensation indeed depends on β-oxidation pathway. We also found that a high ETO dosage induced death in the *Sdhb*^-/-^ mice due to a significant reduction in glucose levels, while no differences were seen in adipose tissue mass (Fig 6F and 6G). Taken together, these results support our notion that HFD can indeed rescue the phenotypes of *Sdhb*^-/-^ mice through the β-oxidation pathway.

**Figure 6.**
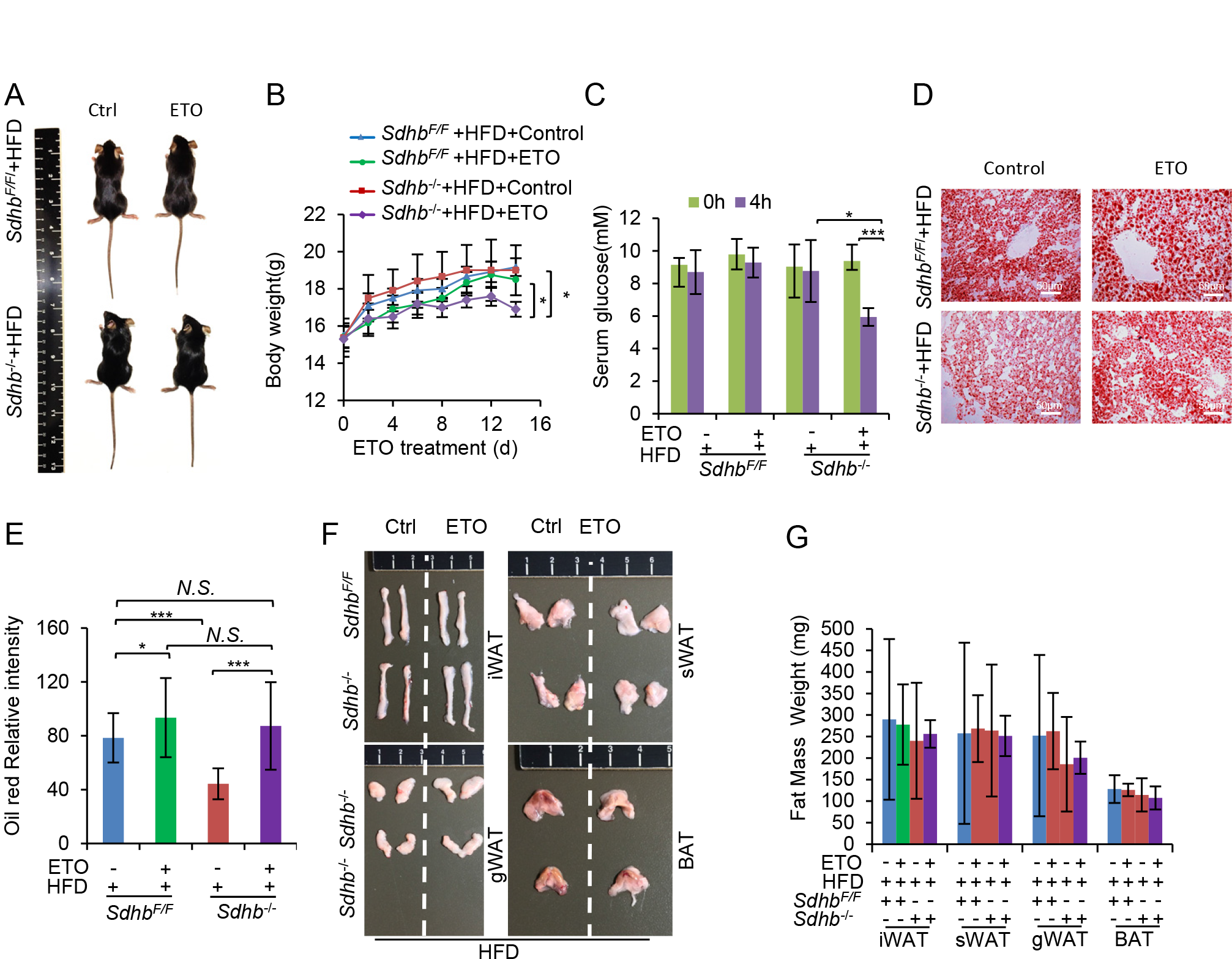
Etomoxir blocks the effect of HFD on *Sdhb^-/-^* mice. A. Body size of ETO-treated mice. *Sdhb^F/F^* and *Sdhb*^-/-^ mice at 4 weeks of age were pretreated with HFD for one week before ETO injection. Animals of each genotype were divided randomly into two groups and given intraperitoneal injections of ETO and olive oil as control, every 2 days for 2 weeks (n=6). B-C. Growth curves (B) and serum glucose levels (C) of ^−^ mice as described in A. D. Representative images showing Oil Red O staining of liver tissues from mice as described in A E. Quantification of the intensity of Oil Red O staining in D (n=4). F. Images of iWAT, sWAT, gWAT and BAT from mice as described in A. G. Quantification of the masses of the main adipose tissues (n=6). Bar graphs represent mean ± SEM, \**P*<0.05, ** *P*<0.01, *** *P*<0.001, *N.S*., no significance.

### Mice with loss function of complex II are resistant to HFD-induced obesity

*Sdhb*^-/-^ mice have accelerated energy expenditure induced by high fatty acid catabolism, so we wondered if they were resistant to HFD-induced obesity. As expected, after treatment with a HFD for almost two years, the *Sdhb*^-/-^ mice were strongly resistant to HFD-induced obesity compared with *Sdhb*^F/F^ mice (Fig 7A). HFD-treated *Sdhb*^F/F^ mice were obese and the body weight was about 2 times more than CD-treated *Sdhb*^F/F^ mice. However, HFD-treated *Sdhb*^-/-^ mice had the same body sizes as CD-treated *Sdhb*^F/F^ mice and lower body weights (Fig 7B and 7C). HFD-treated *Sdhb*^F/F^ mice had higher serum glucose (Fig 7D) and higher body fat ratios than that in CD-treated *Sdhb*^F/F^ mice (Fig 7E), but in HFD-treated *Sdhb*^-/-^ mice, serum glucose and body fat ratios were significantly reduced to the levels observed in CD-treated *Sdhb*^F/F^ mice. The masses of all adipose tissues were also reduced in HFD-treated *Sdhb*^-/-^ mice (Fig 7F and 7G). HFD treatment induced heavy fatty livers in HFD-treated *Sdhb*^F/F^ mice, but no fatty livers syndromes were seen in HFD-treated *Sdhb*^-/-^ mice (Fig 7H and 7I). Lipid accumulation (Fig 7H and 7J) in the livers of HFD-treated *Sdhb*^-/-^ mice was significantly lower than that in HFD- and CD-treated *Sdhb^F/F^* mice. These data indicated that accelerated energy catabolism induced by complex II dysfunction can inhibit fat accumulation in the body, and complex II may therefore be a potential target for combating obesity.

**Figure 7.**
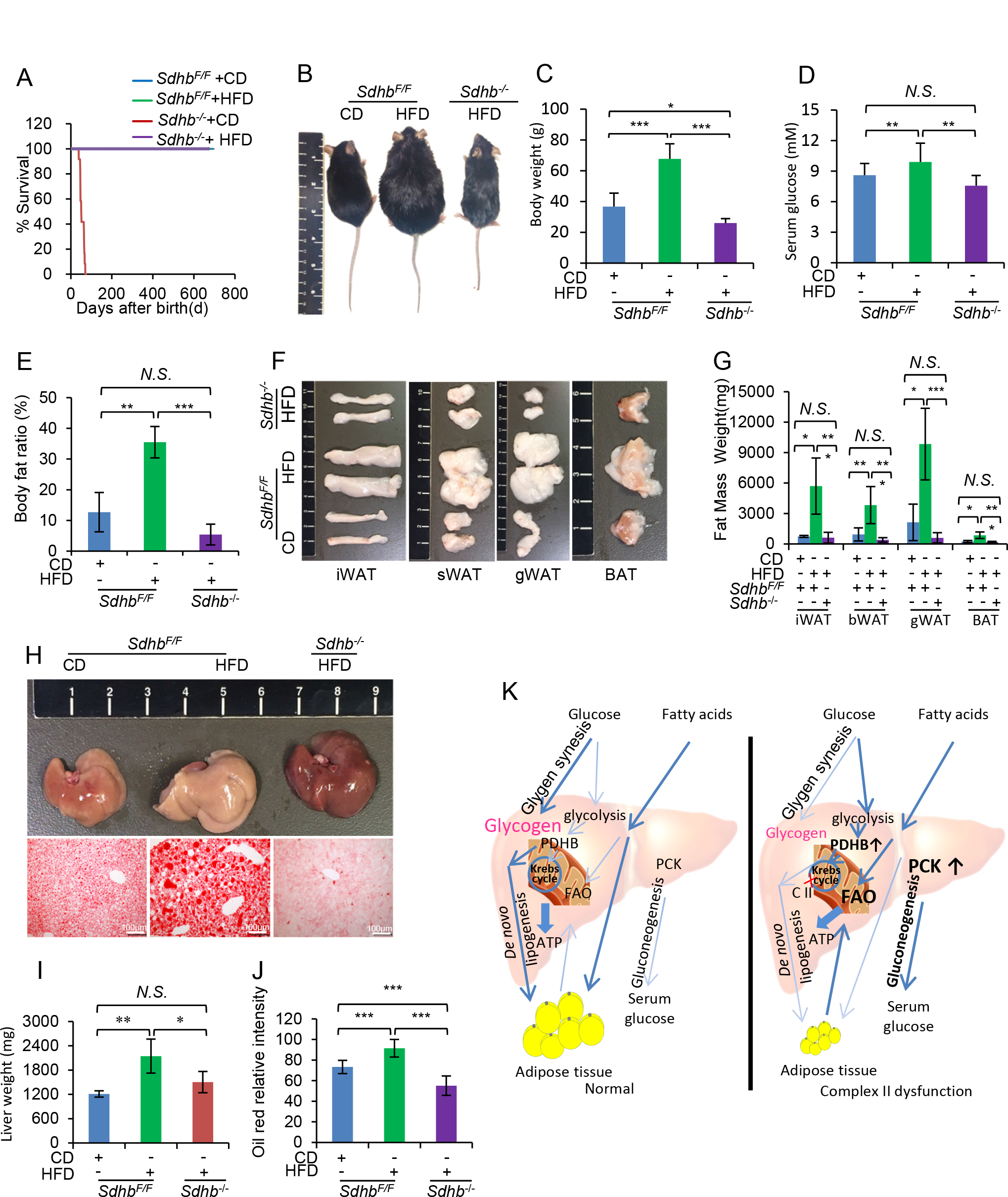
Complex II-deficient mice are resistant to HFD induced-obesity. A. Survival curves of *Sdhb^F/F^* and *Sdhb*^-/-^ mice treated with HFD or CD from 4 weeks of age to two years (n=6-9). B-E. Body sizes (B), body weights (C), serum glucose levels (D) and body fat ratios (E) of mice as described in A. F. Images of iWAT, sWAT, gWAT and BAT tissues of mice as described in A. G. Quantification of the masses of the adipose tissues from F (n=6). H. Images of livers (upper panel) and Oil Red O staining of liver sections (lower panel; scale bar: 50 μm) of mice as described in A. I. Quantification of liver weights in H (upper panel) (n=6). Bar graphs represent mean ± SEM, \**p*<0.05, ** *p*<0.01, *N.S*., no significance. J. Quantification of the intensity of Oil Red O staining in I (lower panel) (n=4; 5 microscope fields were examined per liver section). K. Schematic diagram of complex II regulated energy metabolism and obesity resistance. The results showed dysfunction of Complex II can increase energy expenditure *in vivo*. Supplemented high energy diet could compensate the high energy expenditure. Complex II is a potential target to combat obesity. Bar graphs represent mean ± SEM, \**P*<0.05, ** *P*<0.01, *** *P*<0.001, *N.S*., no significance.

## Discussion

Complex II lies at the crossroads of Krebs cycle and OXPHOS pathways, and is essential for cellular energy metabolism. In the current study, we generated SDHB liver-specific knockout mice and found that SDHB knockout abrogated the assembly and the activity of complex II without affecting other mitochondrial respiration complexes. Loss of complex II reprograms both glucose and fatty acid catabolism and accelerates energy expenditure of substrates, leading to systemic glucose and fat deprivation. These data highlight that mitochondrial respiratory complexes are central regulators of systemic metabolism. Accumulation of succinate, decreases in fumarate and malate, and increased glycolysis were observed in SDHB-mutated cells [23, 24]. Pathways including the ROS-HIF1/2A pathway [25] and the succinate-PHD-HIF1A axis [21] may contribute to these processes. Also, it has been demonstrated that complex II-defective cells increase their reliance on pyruvate carboxylase, which converts pyruvate into the Krebs cycle intermediate oxaloacetate, from which cells can synthesize aspartate, irrespective of the disrupted Krebs cycle [26]. In common with our study, dysfunctions of other respiratory complexes are associated with many diseases. Whole-body knockout of NDUFS4 (a subunit of complex I) can cause multiple disorders and death with encephalomyopathy in mice [27]. Complex I deficiency increases heart failure by altering the redox state [28]. Complex III deficiency induced by UQCRC2 mutation caused recurrent liver failure, lactic acidosis and hypoglycemia in a patient [29]. Loss-of-function mutations in the gene encoding TTC19 (tetratricopeptide repeat domain 19, Complex III) is associated with severe neurological phenotypes [30].

Gene profiling reveals that the loss of complex II reprograms both glucose and fatty acid catabolism, resulting in acceleration of both. Loss of complex II upregulates the expression of genes involved in both glucose and fatty acid catabolism, and the levels of proteins involved in glucose transportation, glycolysis, Krebs cycle and β-oxidation. HIF1A is reported as a transcriptional factor for glucose metabolism and angiogenesis, and undergoes constitutive degradation by a process that involves a series of post-translational modifications [31]. It was shown that inactivation of SDH can induce a pseudohypoxia phenomenon, in which HIF1A is induced under normoxic conditions [20, 21], and the stabilized HIF1A could increase glucose metabolism. Consistent with this theory, cells with complex II deficiency showed increased glucose consumption, including higher glucose uptake and glycolysis [24]. We also found increased HIF1A protein levels and upregulated glucose metabolism in *Sdhb^-/-^* mice. Additionally, we found that Krebs cycle enzymes were upregulated, suggesting that residual enzyme activities are needed for coupling with the cell’s metabolism. Our results are different from those of Kitazawa, who reported that in SDHB-knockout cultured cells there were no changes in protein levels, but there was bidirectional metabolic flow in Krebs cycle to maintain the necessary amounts of metabolites [24]. The discrepancy may be due to the reason that the cellular adaptive response is different between in vivo and in vitro. We were surprised to see the increased levels of proteins in the β-oxidation pathway, which indicated that fatty acids are mobilized as fuels to sustain the energy demand in *Sdhb^-/-^* mice. We also found increased levels of PPARA, which plays a major role in regulating β-oxidation by inducing expression of the gene encoding CPT1 [32]. The increased HIF1A and PPARA activity may underlie the metabolic reprogramming that increases the catabolism of both glucose and fatty acids. Further investigations are needed to clarify the precise pathways involved in these metabolic reprogramming events.

It is striking to find that supplementation with high levels of carbohydrate or fat completely reverses the phenotypes of *Sdhb^-/-^* mice. These dietary supplements restored the serum glucose level and lipid storage in addition to extending the lifespan and blocking body-weight loss. These data clearly showed that lipogenesis from glucose or gluconeogenesis from fatty acids were activated to maintain energy homeostasis. Increased levels of the gluconeogenesis core enzymes PCK1/2 may explain the restored serum glucose levels. These results indicate that complex II not only regulates glucose metabolism, but also fatty acid metabolism, and there exists a mutual compensatory relationship between the glucose and fat metabolism pathways. These findings also highlight that the liver is the metabolic center of the body, and orchestrates the whole body’s energy use of glucose and fatty acids.

Imbalanced energy metabolism causes metabolic syndromes, which include central obesity, insulin resistance, glucose intolerance and non-alcoholic fatty liver disease. Together, these syndromes are among the greatest challenges to human health in the 21^st^-century. Our studies show that there is compensation between energy sources and also between different organs, and offer a unique system to address the complex issue of energy storage and energy expenditure at the organism level. In obese patients, excess body fat is accumulated, which is harmful to cellular functions and causally related with many diseases. Intriguingly, we found that mice with complex II dysfunction are resistant to HFD-induced obesity, which is probably because the loss of complex II accelerates the expenditure of energy derived from both glucose and lipids. Our results thus suggest that complex II is an ideal target for accelerating cellular energy consumption to fight obesity. Future studies are warranted to further investigate the exact mechanisms through which complex II dysfunction regulates glucose and lipid metabolism, and to find new inhibitors that target complex II to fight obesity.

## Acknowledgments

This research was supported by Grants 2016YFA0500201 and 2016YFA0100503 from Ministry of Science and Technology of China to QC, Grants 91754114, 31271529, 31671441 and 301520103904 from the Natural Science Foundation of China to YZ and QC, and Grant QYZDJ-SSW-SMC004 from the CAS Key Project of Frontier Science, and the Beijing Natural Science Foundation (5161002) to QC.

## Author Contributions

CM conceived and designed the experiments with YZ and QC; CM performed most of the experiments and data analysis with assistance from BM, XZ, FG, LC, MW, JL, TZ, HC, QZ, KM, QqL, RC, QL, HW, LL, XW, JW, YgZ, YZ, LiW; CZ performed some biochemical experiments. CM and YZ wrote the manuscript with the help of BM and QC. All authors provided intellectual input and read the manuscript.

## Materials and Methods

### Animals and ethics statement

All mice in this study were housed in the specific-pathogen free (SPF) facility in Nankai University. All germ-free animal diets, chow diet (CD), breeding diet (BD) and high-fat diet (HFD), were purchased from Beijing HFK Bioscience Co., Ltd. Energy supply ratios are listed in expanded view Table 1. All experimental procedures were approved by the Institutional Animal Care of Experimental Animal Center of Nankai University. All efforts were made to minimize suffering and the number of animals used.

**Expanded View Table 1.**
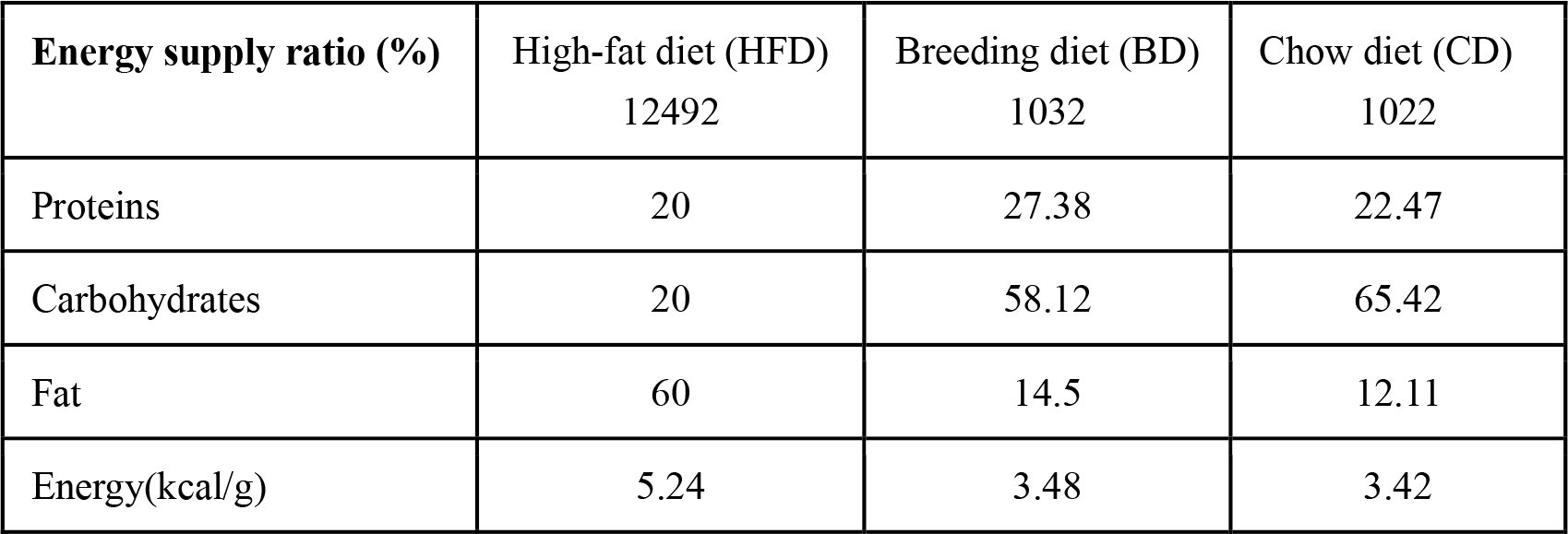
Related to the methods section animals and ethics statement Energy supply ratio of the different mouse diets

**Energy supply ratio of the different mouse diets**
| **Energy supply ratio (%)** | High-fat diet (HFD)<br>12492 | Breeding diet (BD)<br>1032 | Chow diet (CD)<br>1022 |
| --- | --- | --- | --- |
| Proteins | 20 | 27.38 | 22.47 |
| Carbohydrates | 20 | 58.12 | 65.42 |
| Fat | 60 | 14.5 | 12.11 |
| Energy(kcal/g) | 5.24 | 3.48 | 3.42 |

SDHB liver-specific knockout mice were obtained through the Cre-LoxP system. *Sdhb^F/F^* mice were designed in our lab and constructed in the Model Animal Research Center of Nanjing University. *Alb-Cre^+/−^* mice was purchased from the Model Animal Research Center of Nanjing University. In order to breed more *Alb-Cre^+/−^×Sdhb^F/F^* (called *Sdhb^-/-^* for short) mice, we generated 3 strains of mice. Strain 1: *Sdhb^F/F^*, the maintainer strain; strain 2: *Alb-Cre^+/+^ ×Sdhb^F/W^*, the hybrid strain; strain 3: *Alb-Cre^+/+^×Sdhb^W/W^*, for breeding strain 2. Strain 2 and strain 3 were firstly bred through crossing *Alb-Cre^+/−^ ×Sdhb^W/W^* with *Alb-Cre^+/−^ ×Sdhb^F/W^* (offspring of strain 1 and *Alb-Cre^+/−^* mice), and then strain 3 was maintained through selfing, and strain 2 was maintained through crossing strain 3 and strain 2. *Sdhb^-/-^* were obtained by crossing strain 1 and strain 2. Control mice used in the experiments were strain 1, *Sdhb^F/F^* mice.

### Genotyping

3-week-old mice were numbered, then tail samples were boiled in mouse tail lysis buffer (25 mM NaOH and 0.2 mM EDTA (pH 12.0) for 30 minutes and neutralized with 40 mM Tis-HCl (pH 5.0) buffer. The supernatants were collected after centrifuging at 12000 rpm for 5 minutes. 1 μL of supernatant was added as template into a PCR amplification system containing 10 μL 2×Taq-mix PCR buffer (Thermo), 8 μL ddH_2_O, and upstream and downstream primes for *Sdhb* or *Cre*. Primers for genotyping are listed in expanded view Table 2.

**Expanded View Table 2.**
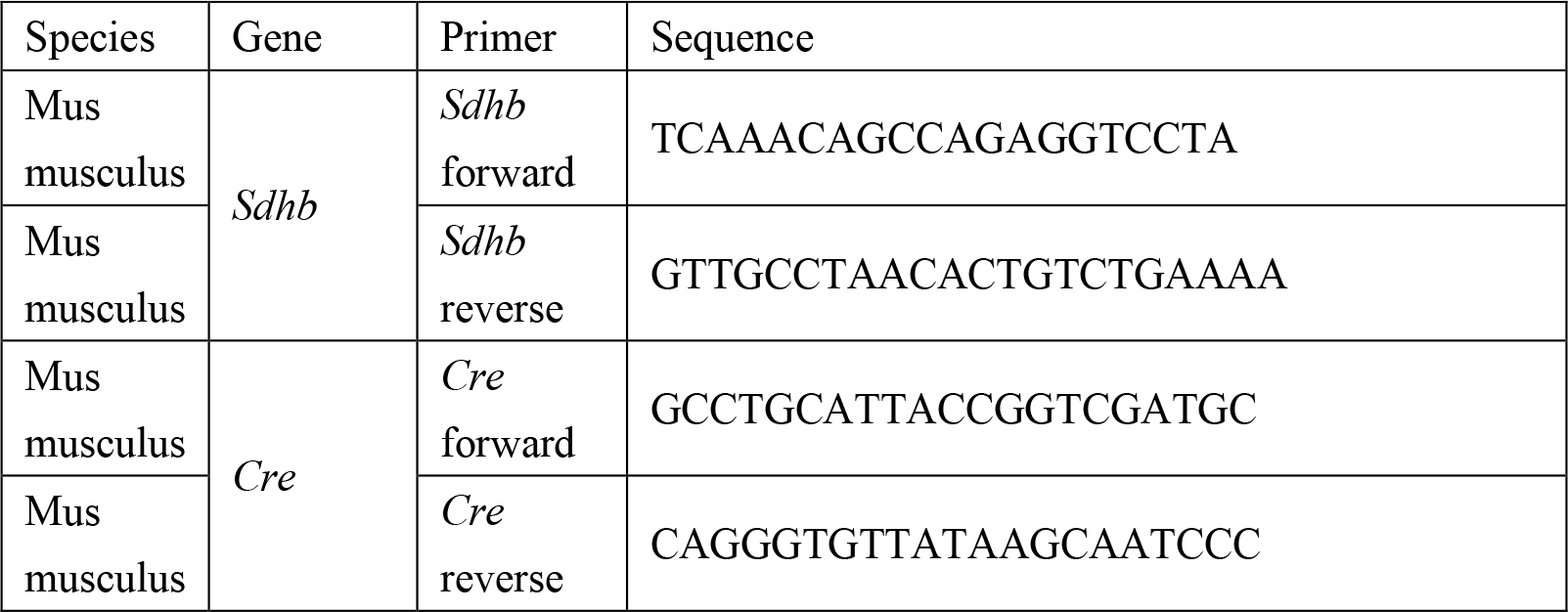
Related to the methods section genotyping Primers used for genotyping

### Animal experiments

One-month-old mice were divided randomly into different groups. For sucrose treatment, mice had free access to CD and water supplemented with or without 20% sucrose. Body weights were measured every 3 days for 2 months. For HFD treatment, mice had free access to HFD and controls had free access to CD. Body weights were measured every 3 days until the end of the experiments. For Etomoxir (ETO) treatment: HFD-treated *Sdhb^F/F^* and *Sdhb*^-/-^ mice were given intraperitoneal injections of Etomoxir (E124862-50, Aladdin) at a dose of 10 mg/kg dissolved in olive oil (BCBQ4885V, Sigma, USA) every 3 days for 3 weeks. Serum glucose was measured at 4 hours after injection, and after recording the viability, the livers of all mice were collected, and fixed or frozen to preserve them.

### Sample collection

Mice were anesthetized and sacrificed by cervical dislocation. All adipose tissue and organs were isolated gently and weighed. Samples were divided into two parts and one part was frozen in liquid nitrogen while the other was fixed in 10% formalin.

### Quantitative RT-PCR

RNA samples were kept in RNAlater buffer, frozen in liquid nitrogen and then stored in a -80 °C freezer. Total mRNA from mouse tissues was extracted with Eastep Super Total RNA Extraction kits (0000278076, Promega). Reverse transcription was performed using M-MLV Reverse Transcriptase (M1705, Promega). Expression of *Sdha, Sdhb, Sdhc, Sdhd* mRNA was analyzed with TransStar Top Green qPCR SuperMix (aq131-04, Trans) and normalized to the house-keeping gene *Actb*. All primers used for RT-qPCR are listed in expanded view Table 3.

**Expanded View Table3.**
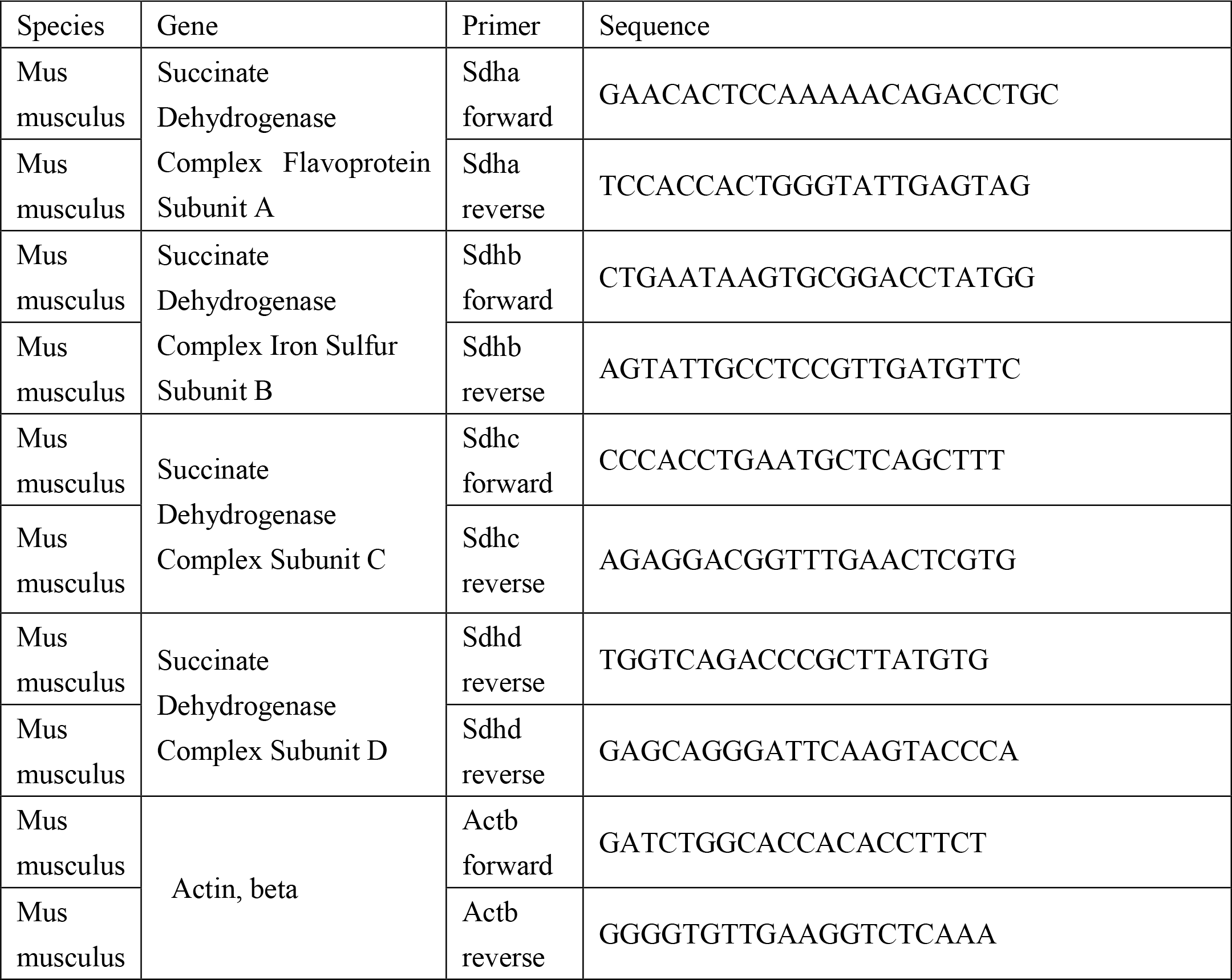
Related to the methods section quantitative RT-PCR Primers used for qPCR

### Isolation of mitochondria and OCR measurement

Anesthetized mice were sacrificed by cervical dislocation, and the liver was removed from the abdominal cavity, 30 mg fresh liver sample were homogenized in a Dounce Tissue Grinder in 1 mL liver mitochondrial isolation buffer I. After centrifuging 2 times at 600 g for 10 min, the supernatant was removed into a new tube and washed 3 times in isolation buffer II (buffer I without BSA) by centrifuging for 10 minutes at 10000 g. Isolated mitochondria were re-suspended in OCR measurement buffer and the concentration was adjusted to 1 mg/mL. After balancing, 20 μL of this mitochondrial suspension was added into the chamber followed by substrates such as succinate or malate and glutamate. ADP was added to initiate the reaction and oxygen consumption was recorded.

### Blue native (BN)-PAGE

BN-PAGE was carried out as previously described [33, 34] and with some modifications. Gradient gels were prepared in our lab. Liver tissue was first homogenized in buffer I to give mitochondria at a concentration of 10 mg/ml, then the mitochondria were lysed in lysis buffer B (50 mM imidazole, 500 mM 6-aminohexanoic acid, EDTA 1 mM pH7.0). n-Decyl-D-maltopyranoside (D000109812, Calbiochem) was added to the mitochondria to give a final concentration of 1.5 mg/mL and the mitochondria were kept on ice for 15 minutes, and then centrifuged at the 13000 rpm for 30 minutes. The supernatant was collected and Coomassie blue was added at a final concentration of 2%. The samples were run on a gel for 30 min at 90 V plus 3 hours at 250 V. Immunoblotting of transmembrane proteins was carried out to detect the assembly of OXPHOS complexes.

### Immunoblots

30 mg frozen liver were homogenized in 1 mL ice-cold RAPA buffer (50 Mm Tris-HCl (pH7.5), 0.15 M NaCl, 1% Na-deoxycholate (W/V), 40 mM EDTA, 0.4% NaF (W/V), 1% Nonidet P40 (V/V), 1 mM PMSF (dissolved in isopropanol at 100 mM and added freshly before use)), centrifuged at 12000 rpm for 10 minutes at 4 °C and the supernatant was collected. 200 μL 6×loading buffer (20 mM Tris-HCl (pH 6.8), 4% SDS, 0.1% bromophenol blue (W/V), 200 mM 2-mercaptoethanol) was added to 1 mL supernatant and the mixture was boiled for 15 minutes. 5 to 10 μL was loaded onto an SDS-PAGE gel running at 70 V and then 110 V to the end. The proteins were transferred to a membrane, and the membrane was blocked in 5% defatted milk. The membrane was incubated with primary and secondary antibodies at 4 °C overnight and room temperature for 2 hours, respectively. Antibodies were used as follows: NDUFA8 (1:1000, Proteintech, 15064-1-AP), SDHA (1:1000, Proteintech, 14865-1-AP), UQCRC1 (1:1000, Proteintech, 21705-1-AP), COXIV (1:1000, Proteintech, 11242-1-AP), ATP5B (1:1000, Proteintech, 17247-1-AP), HIF1A (1:1000, Proteintech, 20960-1-AP), GLUT1¬ (1:1000, Proteintech, 21829-1-AP), GLUT2 (1:1000, Proteintech, 20436-1-AP), GCK (1:1000, Proteintech, 15629-1-AP), ALDOA (1:1000, Proteintech, 11217-1-AP), PDK1 (1:1000, Proteintech, 18262-1-AP), PDHB (1:1000, Proteintech, 14744-1-AP), CS (1:1000, Proteintech, 16131-1-AP), MDH2 (1:1000, Proteintech, 15462-1-AP), ACON2 (1:1000, Proteintech, 11134-1-AP), IDH2 (1:1000, Proteintech, 15932-1-AP), FH (1:1000, Proteintech, 11375-1-AP), MDH2 (1:1000, Proteintech, 15462-1-AP), PPARA (1:1000, Proteintech, 15540-1-AP), PGC1A (1:1000, Proteintech, 20658-1-AP), CPT1A (1:1000, Proteintech, 15184-1-AP), CPT1β (1:1000, Proteintech, 22170-1-AP), CPT2 (1:1000, Proteintech, 26555-1-AP); ACTIN (1:1000, Sigma, A5441); SDHB (1:5000, Abcam, ab14714, 21A11AE7); HADHB (1:2000, Tianjinsaier, SPR09039), ACOX1 (1:1000, Tianjinsaier, SRP06209); antibody: SDHD (1:1000, Cloud-clone Corp., PAK214Mu01).

### SDH activity assay

The DCPIP method was used to measure the activity of complex II with some modifications. Mitochondria were isolated and the concentration was adjusted to 1 mg/mL. A cuvette was filled with 1 mL PBS followed by 40 μL mitochondria, 1 mL of 80 mM DCPIP (BCBC8920V, Sigma, USA), 4 μL of 1 mM Rotenone, and 0.2 μL of 1M ATP. The absorptions at OD 600 were recorded every 1 minute for 5 minutes after initiating the reaction by adding 20 μL of 1 M succinate solution. The DCPIP concentration was calculated with the standard curve formula: C= 0.3229x + 0.0036. The activity of complex II was calculated with the formula: V_SDH_=(C_t_-C_0_)/t/C_mito_.

### Primary hepatocyte isolation

Primary hepatocytes were isolated as previously described with some modifications, as follows. Mice were placed on the dissection board after anaesthetization and a winged needle was inserted into the inferior vena cava after opening the peritoneal cavity. The liver was washed by perfusion with washing buffer (pH 7.4, NaCl 8 g, KCl 0.4 g, NaH_2_PO_4_ 2H_2_O 0.078 g, Na_2_HPO_4_ 12H_2_O 0.151 g, NaHCO_3_ 0.35 g, EGTA 0.019 g, glucose 0.9 g, phenol red 0.006 g per 100 mL) for 20 minutes at a speed of 2 mL/min through the inferior vena cava. The hepatic portal vein was cut after the liver became swollen, and a new perfusion buffer was used (pH 7.4, NaCl 8 g, KCl 0.4 g, NaH_2_PO_4_ 2H_2_O 0.078 g, Na_2_HPO_4_ 12H_2_O 0.151 g, NaHCO_3_ 0.35 g, HEPES 2.38 g, phenol red 0.006 g, 2.5M CaCl_2_ 40μL, collagenase 10 mg per 100 mL) for 10 minutes at the same speed. The liver then became soft and was removed into a dish and hepatocytes were washed with DMEM medium and centrifuged for 3 minutes at 50 g and seeded in 6-well dishes at the concentration of 10^6^ cells/well. Growth medium was DMEM with 10% FBS, penicillin/streptomycin, 100 nM insulin and 100 nM dexamethasone. Growth medium was changed after cell adhesion. Growth medium with or without palmate (PA) was added for the lipid consumption test; After 24 hours, medium was removed, the wells were washed with PBS for three times and the cells were fixed with 10% formalin for oil red staining.

### Histology and chemical stains

For SDH staining, frozen sections of liver samples were incubated in 2.5 mL 0.01M phosphate buffer mixed with 2.5 mL 1M succinate solution and 2.5 mL DMSO containing 5 mg NTB for 15 minutes at room temperature. The sections were washed 3 times with PBS for 3 minutes and fixed in 10% formalin buffer for 5 minutes. The sections were washed again with ddH_2_O before mounting with glycogelatin. Oil Red O staining was carried out as follows: primary isolated hepatocytes in 6-well plates or frozen liver sections were fixed in 10% formalin for 5 minutes, washed 3 times with ddH_2_O for 5 minutes, and then incubated in 60% isopropanol saturated with Oil Red O for 15 minutes. Before the sections were mounted with glycogelatin, they were washed 3 times with ddH_2_O for 5 minutes. Cells were photographed in ddH_2_O. Glycogen was detected with PAS (Periodic Acid-Schiff) staining as follows: paraffin liver sections were deparaffinized and rehydrated with water, then oxidized in 0.8% periodic acid solution containing Buffer I (0.4 g periodic acid (10450-60-9, BBI Life Sciences, China), 35 mL of 95% alcohol, 5 mL of 0.2 M Na2Ac, and 10 mL of ddH_2_O per 50 mL solution). Sections were rinsed 3 times in distilled water for 5 minutes, and then incubated in Schiff reagent (prepared in our lab) for 15 minutes at room temperature. The sections were washed with lukewarm tap water for 5 minutes then with ddH_2_O for 5 minutes before mounting with glycogelatin. Adipose-tissue formalin sections were deparaffinized and rehydrated and stained with Hematoxylin and Eosin (H&E) staining. Sections were dehydrated with gradient alcohol and dimethyl benzene before mounting with neutral balsam. Images were collected using a Leica DM3000 microscope with a Leica DFC420C CCD. Oil Red O and PAS staining images were analyzed by Image J software.

### Glucose tolerance test and glucose measurement

Mice fasted for 4 hours were given intraperitoneal injections of glucose (2 g/kg body weight) for the glucose-tolerance test (GTT). Blood glucose was determined with an ACCU-CHEK glucometer (Roche), at 0, 15, 30, 60, 120 minutes after injection.

### Glycogen accumulation assay

*Sdhb^-/-^* and *Sdhb^F/F^* mice fasted for 24 hours were divided into 3 groups. Intraperitoneal injections of glucose (2 g/kg body weight) were given to groups 2 and 3, while mice in group 1 were given intraperitoneal injections of ddH_2_O. Mice in groups 1, 2, and 3 were sacrificed at 0, 1.5, and 3 hours respectively and liver tissues were fixed in 10% formalin, then embedded in paraffin, sectioned, and stained PAS.

### Statistical analysis

Values shown represent means ± SEM, *p*<0.05 was taken as statistically significant. Data were analyzed using the Student’s t-test. Different symbols in the graphs denote significantly different groups (*p*<0.05). \**p*<0.05; \*\**p*<0.01; \*\*\**p*<0.001, indicate a significant difference from all other groups unless indicated otherwise by lines. *N.S*. =no significance.

## Expanded View Figures

### Figure EV1: Analysis of knock out Sdhb mice

A. Heterozygous whole-body *Sdhb* knockout mice were crossed with *Sdhb^F/F^* to generate *Sdhb* whole-body knockout mice and the pups were genotyped on the first day after birth. The pie chart shows the proportion of pups with each genotype (the numbers are shown in brackets).

B. SDHB protein levels in heart, liver, spleen, lung, kidney, intestine and brain of *Sdhb^F/F^* and liver-specific *Sdhb*^-/-^ mice, GAPDH was used as loading control.

### Figure EV2: Analysis of knock out *Sdhb* mice at 5 weeks of age

A-C. Body sizes (A), body weights (B) and serum glucose levels (C) of *Sdhb^F/F^* and *Sdhb*^-/-^ mice at 5 weeks of age (n=11).

D. PAS staining for glycogen in liver tissues of *Sdhb^F/F^* and *Sdhb^-/-^* mice. Scale bar: 50 μm.

E. GTT results in *Sdhb^F/F^* and *Sdhb*^-/-^ mice at 5 weeks of age (n=7).

F. Body fat ratios of *Sdhb^F/F^* and *Sdhb*^-/-^ mice at 5 weeks of age (n=7).

G. Images of adipose tissues (iWAT, sWAT, gWAT and BAT) from *Sdhb^F/F^* and *Sdhb*^-/-^ mice at 5 weeks of age.

H. Quantification of the adipose tissue mass from G (n=7).

I. H&E staining of paraffin sections of iWAT, sWAT, gWAT and BAT tissue from *Sdhb^F/F^* and *Sdhb*^-/-^ mice at 5 weeks of age (n=4). Scale bar: 50 μm.

J. Oil Red O staining of lipid droplets in the liver tissue of *Sdhb^F/F^* and *Sdhb*^-/-^ mice at 5 weeks of age (n=4). Scale bar: 50 μm.

K. Protein levels of HIF1A, the glucose transporter GLUT1, and enzymes involved in glycolysis (GCK, ALDOA, PKLR, PDK1 and PDHB), Krebs cycle (ACON2 and MDH2), lipid metabolism (CPT2, HADHB and ACOX1) and gluconeogenesis (PCK1) in *Sdhb^F/F^* and *Sdhb*^-/-^ mice at 5 weeks of age. ACTIN was the loading control (n=3).

Bar graphs represent mean ± SEM, \**P*<0.05.

**Abbreviation**
NDUFA8: NADH: Ubiquinone Oxidoreductase Subunit A8.
SDHA: Succinate Dehydrogenase Complex Flavoprotein Subunit A.
SDHB: Succinate Dehydrogenase Complex Iron Sulfur Subunit B.
UQCRC1: Ubiquinol-cytochrome c reductase core protein I.
CoxIV: cytochrome c oxidase subunit IV isoform 1.
ATP5B: ATP Synthase, H+ Transporting, Mitochondrial F1 Complex, Beta Polypeptide.
Glut1: Glucose Transporter Type 1.
Glut2: Glucose Transporter Type 1.
GCK: Glucokinase.
ALDOA: Aldolase, Fructose-Bisphosphate A.
PDK1: Pyruvate Dehydrogenase Kinase 1.
PDHB: Pyruvate Dehydrogenase E1 Beta Subunit.
PCK1: Phosphoenolpyruvate Carboxykinase 1.
PCK2: Phosphoenolpyruvate Carboxykinase 2, Mitochondrial.
HIF1A: Hypoxia Inducible Factor 1 Alpha Subunit.
CS: Citrate Synthase.
Acon2: Aconitase 2.
IDH2: Isocitrate Dehydrogenase (NADP^+^) 2, Mitochondrial.
GAPDH: Glyceraldehyde-3-Phosphate Dehydrogenase.
OGDH: Oxoglutarate Dehydrogenase.
SDHD: Succinate Dehydrogenase Complex Subunit D.
FH: Fumarate Hydratase.
MDH2: Malate Dehydrogenase 2.
PPARA: Peroxisome Proliferator Activated Receptor Alpha.
PGC1A: PPARG Coactivator 1 Alpha.
CPT1A: Carnitine Palmitoyltransferase 1A.
CPT1B: Carnitine Palmitoyltransferase 1B.
CPT2: Carnitine Palmitoyltransferase 2.
HADHB: Hydroxyacyl-CoA Dehydrogenase/3-Ketoacyl-CoA Thiolase/Enoyl-CoA Hydratase (Trifunctional Protein), Beta Subunit.
ACOX1: Acyl-CoA Oxidase 1.
OXPHOS: Oxidative phosphorylation.
HFD: high fat diet.
CD: chow diet.
BD: breeding diet.
VLDL: very low-density lipoproteins.

